# Fibronectin Coating of Tissue Culture Polystyrene to Improve Superficial Zone Chondrocyte Expansion

**DOI:** 10.64898/2026.07.02.736120

**Authors:** John E. Caputo, Thomas J. Manzoni, Isabella Ewine, Alvin W. Su, Justin Parreno

**Author notes:** Correspondence to: Justin Parreno, Wolf Hall, 105 The Green, Newark, Delaware, 19716, USA. Denotes Co-First Author (John E. Caputo, Thomas J. Manzoni).

## Abstract

The surface layer of articular cartilage provides for low-friction joint movement and protects the tissue from mechanical wear. The superficial zone chondrocytes (SZCs) of the surface layer produce proteoglycan-4 (PRG4), which is a lubricant that is necessary to reduce friction. Articular cartilage has limited capacity for self-repair and cell-based therapies, such as autologous chondrocyte implantation (ACI), is used to stimulate repair. However, in ACI, cells are expanded on tissue culture polystyrene where SZC poorly attach, proliferate slowly and dedifferentiate. Consequently, expanded SZC produce fibrocartilage tissue with insufficient PRG4. We previously demonstrated that culturing SZC on chondrocyte-derived decellularized extracellular matrix (CM) enhances SZC attachment and preserves phenotype. Since fibronectin (FN) was identified as the most abundant matrix protein within CM, here we tested the hypothesis that FN-coated culture surfaces would partially reproduce the beneficial effects of CM. We found that, similar to CM, SZC on FN-coated polystyrene increased SZC attachment and proliferation. However, unlike CM, SZCs expanded on FN-coated polystyrene remained more dedifferentiated as indicated by spread cells, elevated fibroblastic and contractile mRNA levels, and increased formation of αSMA positive stress fibers. Consistent with the dedifferentiated phenotype, SZC on FN-coated polystyrene displayed extensive stress fibers, and higher nuclear myocardin-related-transcription-factor-a (MRTF-A). In contrast, CM reduced stress fiber formation and diminished nuclear MRTF-A in SZC. CM provides matrix cues beyond FN that suppress dedifferentiation and preserve the SZC phenotype. Identifying the matrix cues necessary to improve SZC expansion could lead to the generation of a superior surface in ACI repair tissue.

## INTRODUCTION

Articular cartilage functions to provide a smooth and lubricated surface, facilitating load distribution during joint articulation. The bulk of articular cartilage matrix is made up of primarily type II collagen (COL2) and aggrecan (ACAN). The matrix of cartilage is further organized into three distinct zones, each with unique structures and functions [1, 2]. The top surface layer, or superficial zone (SZ), plays a crucial role in providing a lubricated surface for joint movement. Superficial zone chondrocytes (SZCs) are essential for maintaining SZ function by producing lubricating proteins such as proteoglycan-4 (PRG4) to maintain frictionless movement [3, 4]. Cartilage is incapable of self-repair after injury and as a result, cartilage injuries present significant challenges [2]. Cell-based therapies such as autologous chondrocyte implantation aim to restore the damaged tissue by expanding chondrocytes for cell number *in vitro* and implanting them into the defect site [5–8]. While these approaches can aid tissue repair, their long-term success is limited by the challenges associated with chondrocytes expansion.

A major challenge is the production of an adequate superficial zone due to several underlying issues. SZCs exhibit reduced attachment and slower proliferation than deeper-zone chondrocytes during monolayer expansion, resulting in their progressive depletion from the expanded chondrocyte population [9, 10]. The importance of maintaining SZCs is further highlighted by the fact that the superficial zone is among the earliest regions affected during cartilage degeneration, making preservation of SZCs a critical challenge for cartilage regeneration strategies [2]. In addition, the expansion of SZCs on monolayer culture alters the SZ phenotype, where loss of SZC function during expansion is linked to chondrocyte dedifferentiation [10]. On polystyrene (PS), SZCs undergo alterations in morphology characterized by increased cell spreading and elongation. These changes coincide with reduced expression of chondrogenic molecules such as COL2, ACAN and PRG4 as well as increased expression of fibroblastic matrix molecules like type I collagen (COL1) and tenascin-C (TNC) [11–15]. The increased production of fibroblastic matrix molecules contributes to the formation of mechanically inferior fibrocartilage rather than native hyaline cartilage [16, 17]. In addition, monolayer expansion results in increased expression of contractile markers like alpha smooth muscle actin (αSMA), an isoform of actin that contributes to the highly contractile phenotype in dedifferentiated cells [11–15, 18]. Increased contractility could lead to tissue contraction of regenerated tissue and impair integration with surrounding native tissue following implantation.

The reorganization of the actin cytoskeleton is a key regulator of chondrocyte phenotype and contributes to the progression of dedifferentiation during monolayer expansion [11, 12, 19, 20]. This reorganization is characterized by a shift in the proportion of globular (G-) actin to filamentous (F-) actin within cells. In monolayer culture, this increased polymerization of G-actin into F-actin promotes the formation of stress fibers, leading to the spread of fibroblastic morphology [12, 21]. As actin is an important regulator of the chondrocyte phenotype, disrupting F-actin stress fibers suppresses the dedifferentiated phenotype in chondrocytes [19, 22]. Alterations in G/F-actin regulates downstream transcription through regulation of G-actin binding myocardin-related transcription factor-a (MRTF-A). When G-actin levels are low, MRTF-A is predominantly nuclear where it promotes fibroblastic gene expression. Conversely, when G-actin levels are high, MRTF-A binds to G-actin and remains in the cytoplasm limiting transcription [23, 24]. While MRTF-A has been shown to regulate phenotype in primary SZC chondrocytes, its role during the expansion of SZC has not yet been defined [25].

Strategies that improve SZC expansion and suppress dedifferentiation of SZC by targeting key pathways, such as actin-MRTF-A signaling, are needed to overcome the limitations of current cartilage repair approaches. Monolayer culture of SZC on chondrocyte-derived decellularized extracellular matrices (CM) improves SZC culture by increasing primary cellular attachment, increasing proliferation, and limiting the increase in fibroblastic and contractile gene expression during SZC expansion [26]. These effects address key limitations in SZC expansion of low SZC yield and excessive dedifferentiation during culture. However, the underlying matrix molecules and mechanisms that improve SZC attachment and phenotype on CM remain unclear. Our analysis of CM matrix content identified fibronectin (FN) as the most abundant protein present on CM [26]. FN regulates SZC adhesion, proliferation and differentiation status through interactions with cell surface integrins, and SZCs preferentially bind FN due to their high expression of the fibronectin-binding integrin, α5β1 [27, 28]. While cell attachment may be enhanced by coating culture surfaces with FN, FN signaling through integrins has been shown to regulate actin polymerization and cytoskeletal organization, processes associated with MRTF-A activation and chondrocyte dedifferentiation [29]. Therefore, while FN may improve SZC attachment and proliferation it could also promote critical signaling pathways associated with dedifferentiation. Therefore, it remains unclear whether FN alone can reproduce the beneficial effects of CM on SZC expansion while maintaining the SZC phenotype.

In this study, we hypothesize that coating PS with FN will improve SZC attachment and proliferation in 2D culture. However, since FN has been implicated in actin polymerization and MRTF-A activation, we hypothesize that FN will also promote SZC dedifferentiation during monolayer expansion. To test this, SZCs will be cultured and expanded on PS, FN-coated PS, or CM-coated PS. We will assess cell attachment, expansion rate, F-actin organization and MRTF-A nuclear activity, and expression of chondrogenic and dedifferentiation molecules. Together, this study aims to define whether FN improves monolayer expansion of SZCs that could be subsequently used for tissue regeneration purposes.

## RESULTS

### Culture of P0 SZC on CM and FN Enhance Primary SZC Attachment

To determine whether FN enhances primary SZC attachment, we seeded equivalent numbers of P0 SZCs on PS, CM, and FN. We found that after 24 hours, seeding SZCs on CM and FN coatings increases the percent area covered by cells (Figure 1A). SZCs cultured on CM and FN exhibit 8.4% ± 2.7% and 9.9% ± 3.1% coverage, respectively, compared to 3.5% ± 0.9% on PS (Figure 1B). We also measured the proportion of cells that attached after 24 hours of culture using DNA quantification. On CM and FN, SZCs attachment is 32.4 ± 4.7% and 32.5 ± 7.6% respectively (Figure 1C). Attachment is greater on CM and FN then PS as cell attachment on PS is 16.8 ± 3.8%. These findings indicate that CM and FN enhance primary SZC attachment relative to PS.

**Figure 1.**
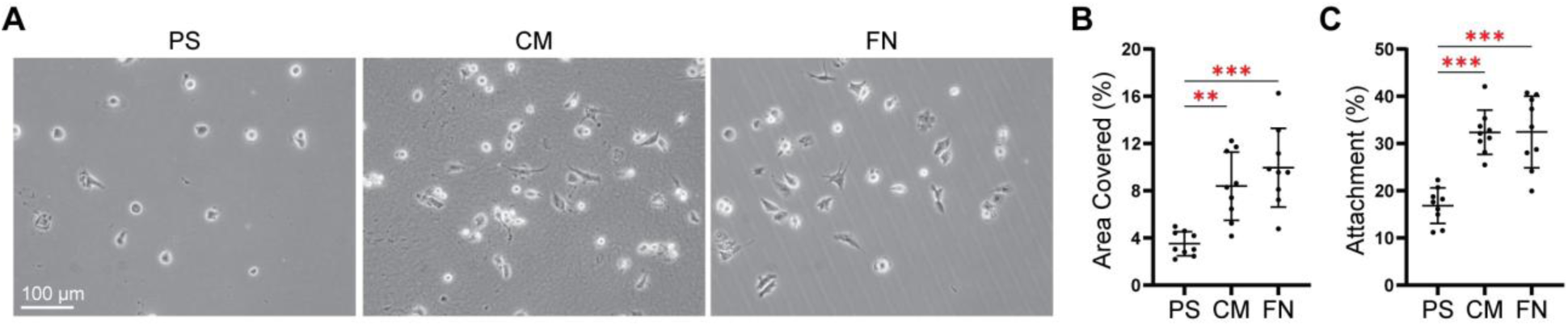
Attachment of primary SZC on PS, CM, and FN. (A) Light microscopy images and quantified (B) % area covered of primary (P0) SZC attached to PS, CM, and FN after 24 hours. (C) Percent attachment of SZC seeded on PS, CM, and FN after 24 hours. **p < 0.01 ***p < 0.001.

### Expansion of SZC on CM and FN Reduce Time to Confluency

Next, we measured the length of time required for SZCs in monolayer culture to reach confluency (80-90% cell coverage). Culturing SZC on CM and FN reduce the time required to reach confluency during expansion from P0 to P1. SZCs cultured on CM and FN reach confluency on day 8, whereas SZCs cultured on PS reach confluency on day 12. On day 8, when SZCs on CM and FN reach confluency, SZCs cultured on PS achieve only 37.9% ± 5.6% confluency (Figure 2A and B).

**Figure 2.**
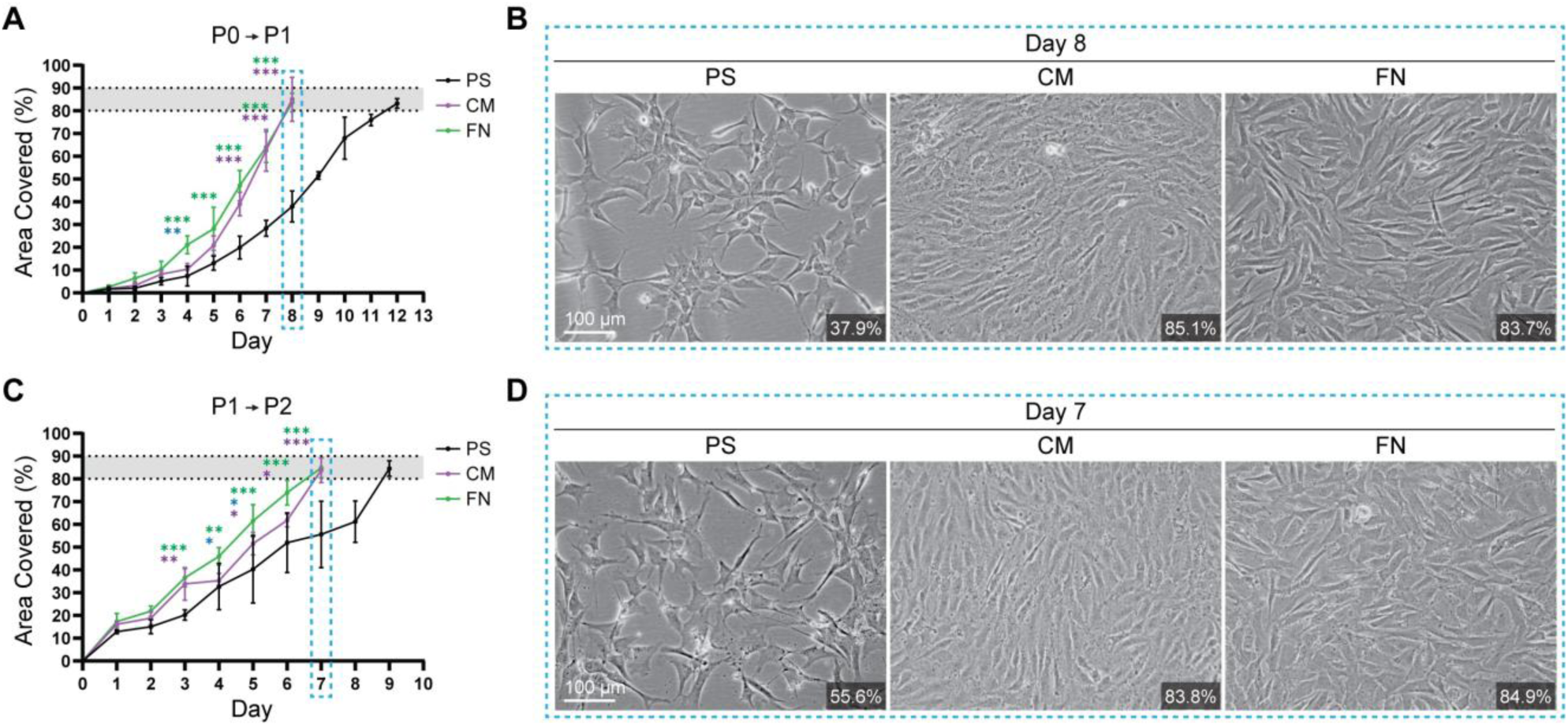
Confluency of SZC cultured on PS, CM, and FN from (A, B) P0 to P1 and (C, D) P1 to P2. (A, C) % confluency of passaged SZC on PS, CM, or FN each day as cells progress from (A, B) P0 to P1 and (C, D) P1 to P2. (B, D) Representative light microscopy images at Day 8 and Day 7, respectively. *p<0.05, **p<0.01, ***p< 0.001. Comparisons between groups are denoted by color (Green: PS vs FN; Magenta: PS vs CM, Blue: CM vs FN).

Similarly, culturing SZC on CM and FN reduces the time required for SZCs to reach confluency during expansion from P1 to P2. SZCs cultured on CM and FN reach confluency on day 7, compared to day 9 on PS. On day 7, when CM and FN reach confluency, SZCs on PS achieve only 55.6% ± 11.9% confluency (Figure 2C and D). Collectively, these results indicate that CM and FN reduce the time for SZCs to reach confluency compared to PS.

### Passaging SZC on FN Leads to Fibroblastic Morphology and Elongation

We next sought to determine the extent to which expansion on PS, CM and FN influences cell morphology, as an increase in chondrocyte size is associated with dedifferentiation [11]. At P0, SZCs cultured on PS, CM and FN showed no significant differences in cell area or circularity (Figure 3A). Expansion from P0 to P2 increases cell area in SZCs cultured on all substrates. By P2, SZCs on FN and PS are larger (Figure 3A and B) as compared to SZCs cultured on CM.

**Figure 3.**
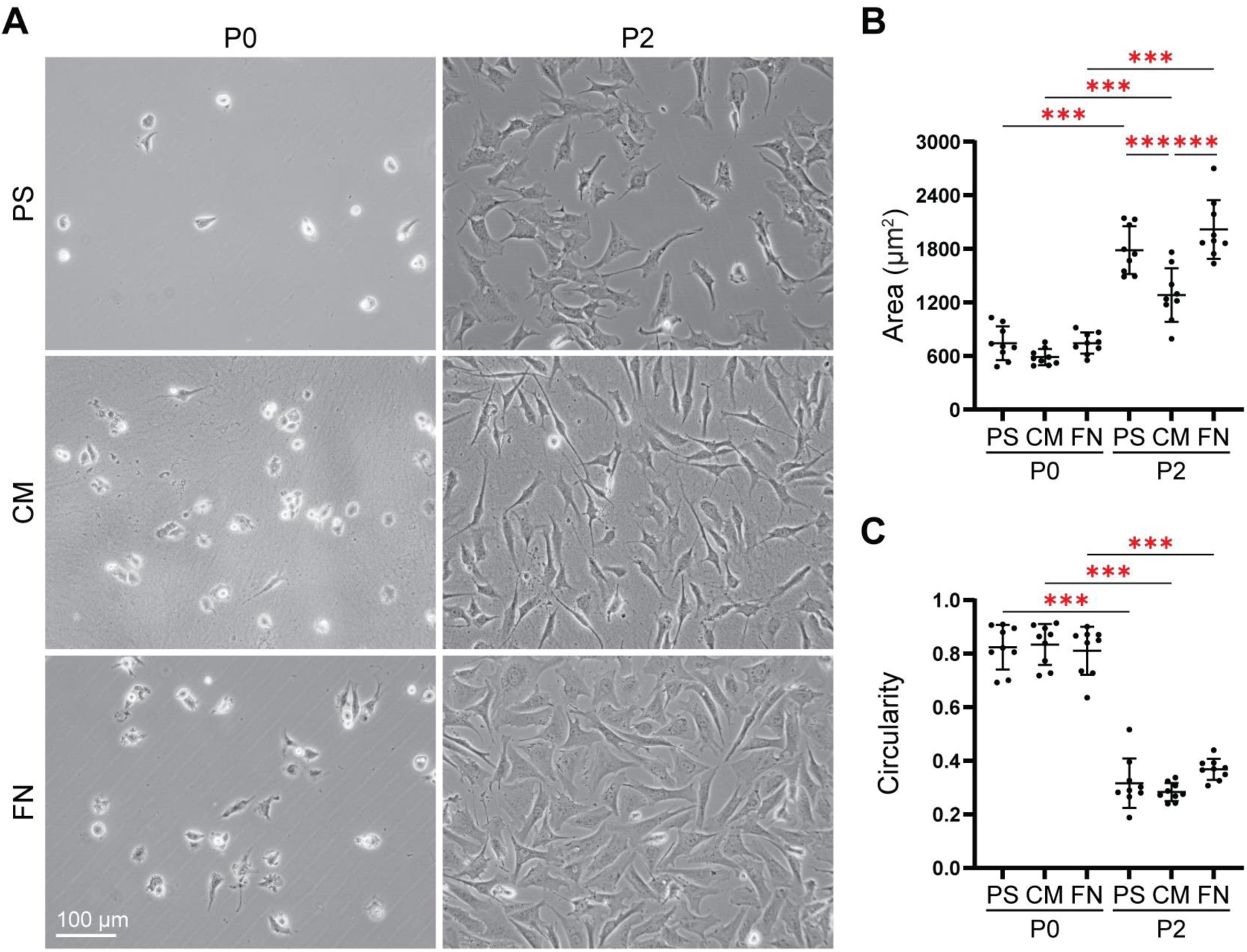
Morphological analysis of P0 and P2 SZC on PS, CM, and FN. (A) Representative light microscopy images of SZC cultured on PS, CM, and FN at P0 and passage 2 P2. Quantification of cell (B) area and (C) circularity at both P0 and P2. ***p < 0.001.

Analysis of cell circularity showed that passaging reduces cell circularity across all conditions (Figure 3C). Notably, culturing SZCs on FN resembles the morphological changes similar to those seen on PS. Collectively, these results show that CM limits the increase in cell size associated with passaging, whereas FN promotes cell spreading and enlargement similar to PS.

### Expansion of SZC on FN Does Not Prevent Loss of Chondrogenic and SZ-Specific Gene Expression

To evaluate the impact of monolayer expansion of SZC culture on either PS, CM or FN at each passage on SZC dedifferentiation, we measured the mRNA levels of expanded SZCs. For chondrogenic expression (Figure 4A), *ACAN* mRNA levels are similar across PS, CM and FN at P0. Across all culture conditions, *ACAN* mRNA levels decreased significantly from P0 to P2. At P2, there are no significant differences between culture surfaces at this passage. Across all surfaces, *COL2* mRNA levels decrease from P0 to P2. At P0 and P2, *COL2* mRNA levels are similar between SZCs on PS, CM and FN. SRY-Box Transcription Factor 9 (*SOX9*) is a known transcriptional activator of *ACAN* and *COL2* [30, 31]. We found *SOX9* mRNA levels are higher in P0 SZCs on FN as compared to P2 SZCs on FN. We found there are no differences between P0 SZCs on PS, CM, and FN. Similarly, there are no differences between P2 SZC on PS, CM, and FN.

**Figure 4.**
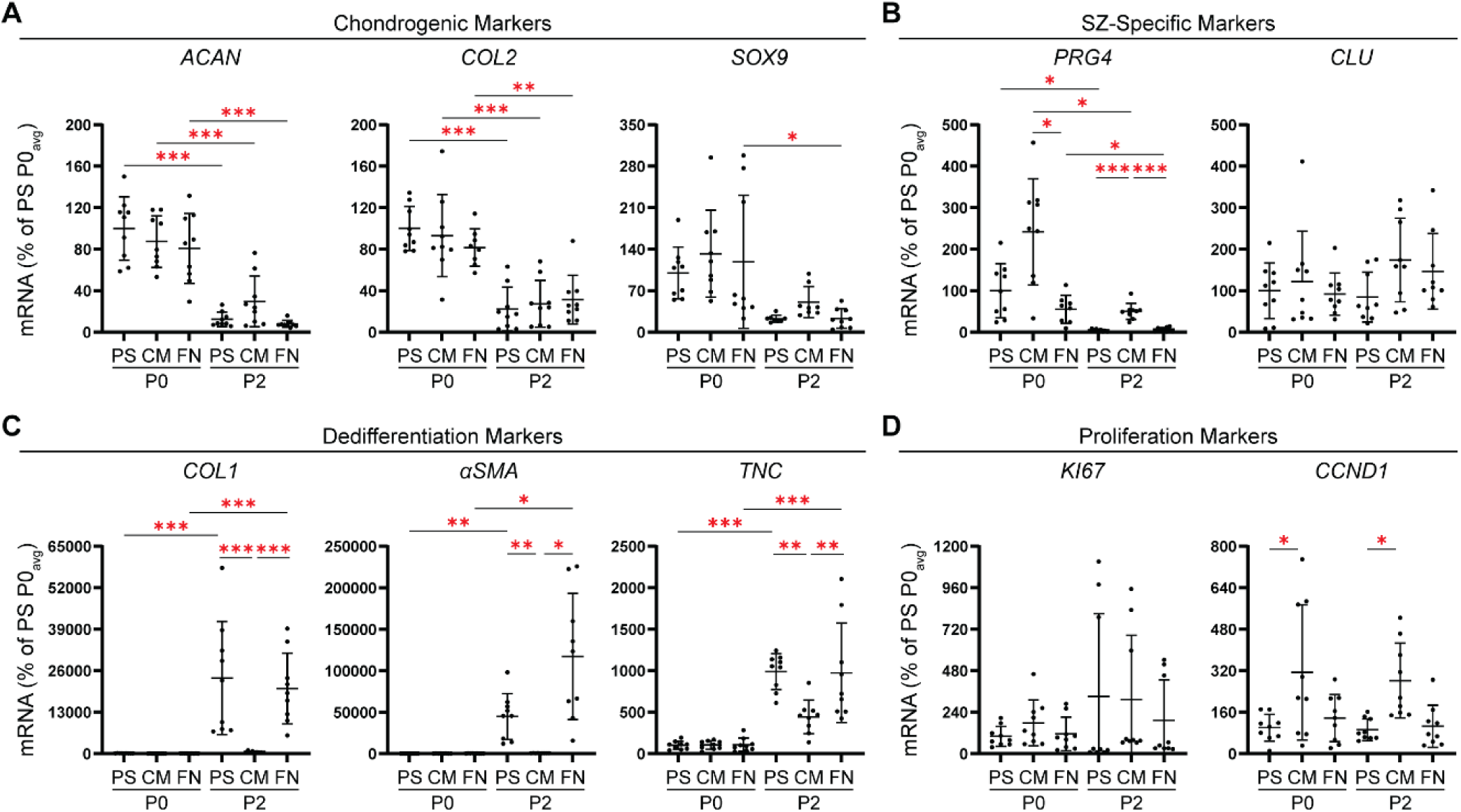
Gene expression analysis of SZC passaged on PS, CM, and FN at P0 and P2. Relative RT-PCR of (A) chondrogenic (*ACAN*, *COL2*, and *SOX9*), (B) SZ-specific (*PRG4* and *CLU*), (C) dedifferentiation (*COL1*, *αSMA*, and *TNC*), and (D) proliferation (*KI67* and *CCND1*) mRNA levels of SZC passaged on PS, CD-ECM (CM), and FN at P0 and P2. Expressed as a percent of P0 PS. *P < 0.05, **P < 0.01, ***P < 0.001.

SZ-specific *PRG4* mRNA levels (Figure 4B) are higher at P0 for SZCs seeded on CM compared to those on FN and PS. Although *PRG4* mRNA levels decrease with passaging across all conditions, SZCs cultured on CM to P2 maintain higher *PRG4* expression than those cultured on PS and FN. There are no differences in SZ molecule *CLU* mRNA levels across all conditions. To evaluate the acquisition of fibroblastic characteristics throughout passaging on the different coatings, we examined *COL1, ɑSMA*, and *TNC* mRNA levels (Figure 4C). Throughout passaging, *COL1, TNC,* and *ɑSMA* mRNA levels increase between P0 and P2 in SZCs cultured on PS and FN, while their expression remains similar between P0 and P2 in SZCs cultured on CM. At P2, *COL1, TNC,* and *ɑSMA* mRNA levels are significantly lower on CM, compared to PS and FN.

Since SZCs passaged on CM and FN reach confluency faster than SZCs on PS, we also examined the mRNA levels of the marker of proliferation Kiel-67 (*KI67)* and the cell cycle regulator cyclin D1 (*CCND1*) (Figure 4D). *KI67* mRNA levels showed substantial variability and did not differ significantly among each culture condition at either passage (P0 and P2). *CCND1* mRNA levels are higher in SZCs on CM relative to PS at both P0 and P2, whereas SZCs cultured on FN show no significant differences from PS. Collectively, these findings indicate that expansion of SZCs on CM, but not FN, represses dedifferentiation-associated mRNA expression levels during passaging.

### Expansion of SZC on FN Does Not Prevent Cytoskeletal Remodeling Associated with Dedifferentiation

αSMA is highly expressed in dedifferentiated chondrocytes and contributes to the highly contractile phenotype of P2 chondrocytes [12, 21] Here, we used αSMA as an indicator of the dedifferentiated phenotype by examining αSMA organization in P2 SZCs cultured on PS, CM and FN (Figure 5). Confocal imaging revealed that 22.0% ± 3.6% of SZCs displayed αSMA positive stress fibers on PS. The proportion of SZCs with αSMA positive stress fibers is similar to SZCs on FN where 18.7% ± 2.9% display αSMA positive stress fibers. In contrast, SZCs cultured on CM display less as only 4.6% ± 2.6% SZCs display αSMA positive stress fibers.

**Figure 5.**
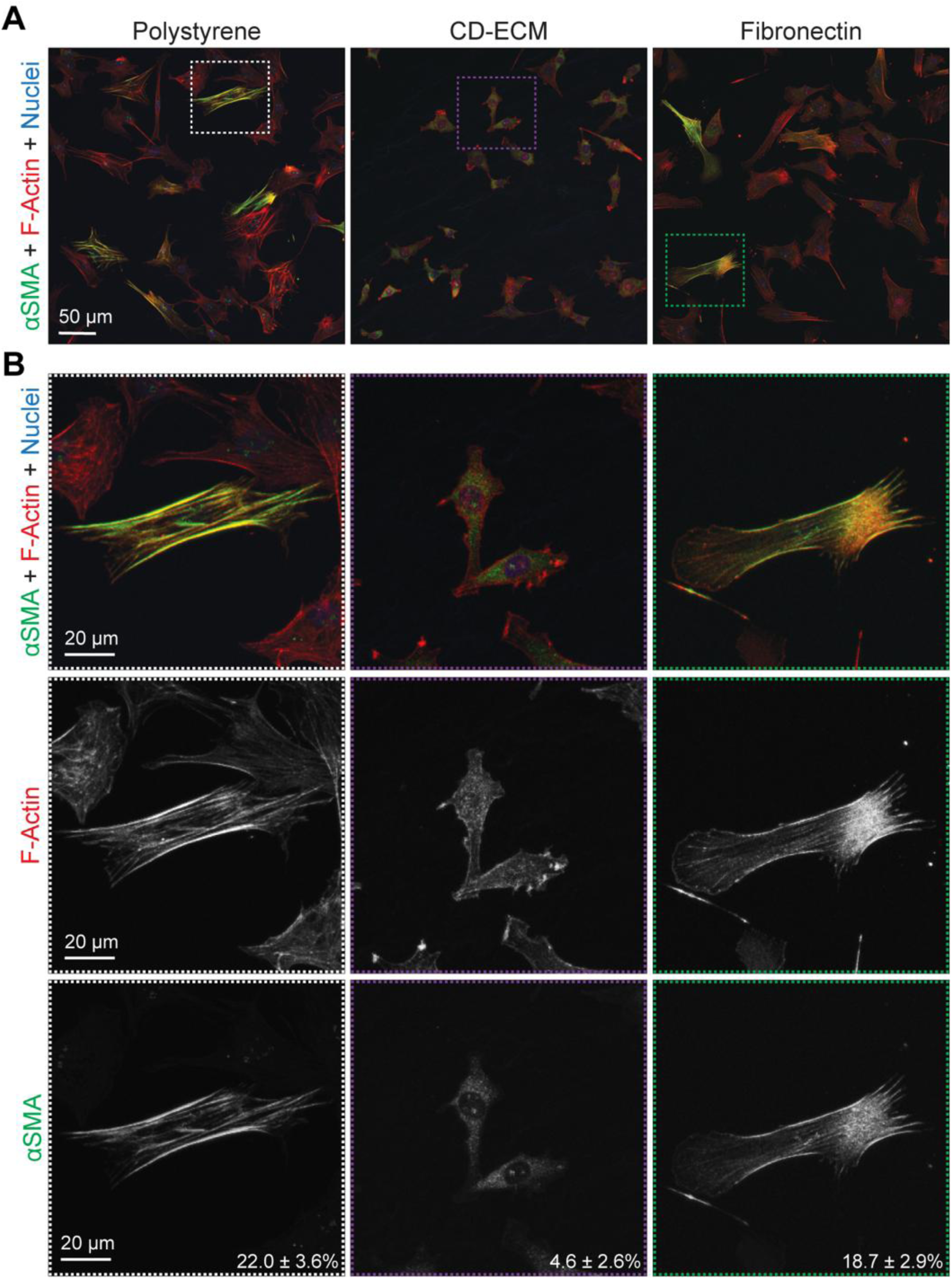
αSMA staining of P2 SZC on PS, CM, and FN. **(A)** Confocal microscopy images showing αSMA (Green), F-Actin (Red), and Nuclei (Blue) in P2 SZC cultured on PS, CM, and FN. Dashed boxes indicate representative cells shown at higher magnification in (B). **(B)** Representative high magnification images of αSMA and F-actin organization, including individual αSMA and F-actin channels. Values indicate the % of P2 SZC with αSMA positive stress fibers on PS, CM, and FN.

### FN Expansion is Associated with Increased MRTF-A Nuclear Localization and Actin Polymerization

Since alteration to the polymerization status of chondrocytes regulates MRTF-A localization to regulate dedifferentiation, [12, 21] we evaluated F-actin organization and MRTF-A localization in P2 SZCs in PS, CM and FN (Figure 6). SZCs cultured on PS and FN display prominent stress fibers as compared to SZCs on CM where there are less F-actin stress fibers (Figure 5B; middle panel and Figure 6A; top panel). In addition, CM reduces MRTF-A nuclear localization in P2 SZCs (Figure 6A; bottom panel, Figure 6B). These results indicate that FN allows for prominent F-actin stress fiber formation and nuclear MRTF-A localization, both of which are associated with SZC dedifferentiation. On the other hand, CM represses F-actin stress fibers and nuclear MRTF-A leading to preservation of the SZC phenotype.

**Figure 6.**
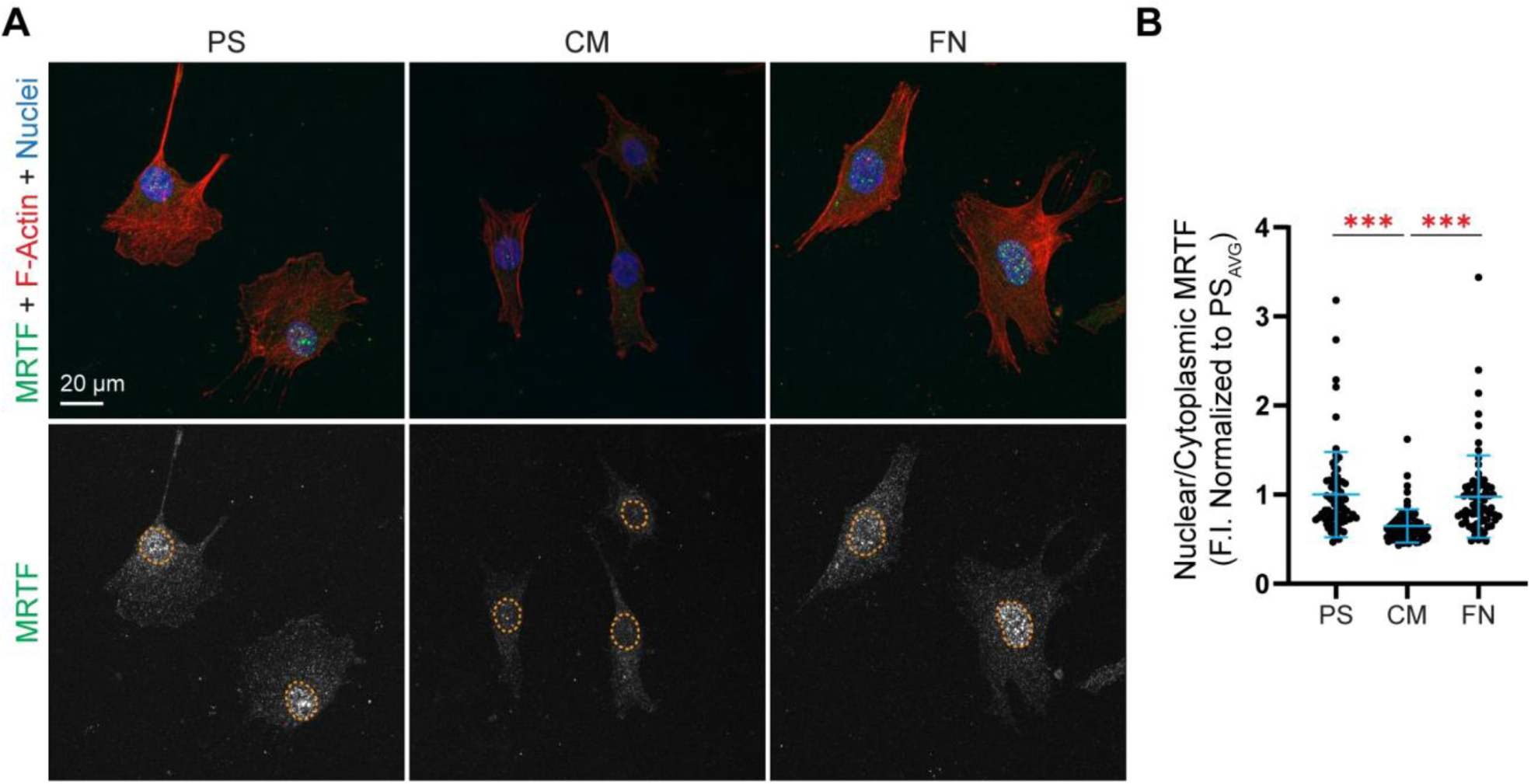
MRTF-A staining and localization in P2 SZC on PS, CM, and FN. **(A)** Confocal microscopy images showing MRTF-A (Green), F-Actin (Red), and Nuclei (Blue) and **(B)** quantification of MRTF-A nuclear and cytoplasmic mean fluorescence intensity of P2 SZC on PS, CM, and FN. ***p < 0.001.

## DISCUSSION

The findings of this study supported our hypothesis that FN could partially reproduce the beneficial effects of CM during SZC expansion. FN increased primary SZC attachment and reduced time to confluency, consistent with our prediction that FN will improve expansion for cell number. However, unlike SZCs expanded on CM, the culture of SZCs on FN did not repress the dedifferentiated phenotype. Instead, SZCs expanded on FN had morphological, transcriptional, and F-actin organization that resemble dedifferentiated SZCs cultured on PS. Therefore, while FN improves SZC attachment and confluency, FN alone is not sufficient to preserve the SZ phenotype during 2D expansion. These findings suggest that FN supports attachment and growth, but additional factors provided by the CM repress the dedifferentiation that occurs during monolayer expansion.

The improved attachment seen on FN is likely mediated through interactions between SZCs and FN-binding integrins. Chondrocytes express several integrins that mediate adhesion to matrix ligands, including receptors that bind FN, collagens, laminin, and osteopontin [27]. Among these, the α5β1 integrin is a well-characterized receptor for FN and has been reported to be highly expressed in SZCs [32, 33]. Previous studies have shown that SZCs preferentially bind to FN substrates, suggesting that α5β1-mediated adhesion may contribute to the improved attachment on FN-coated surfaces [28, 34]. SZCs cultured attach to CM and FN to a similar extent, suggesting that FN may contribute to the improved initial attachment observed on CM.

SZCs cultured on CM and FN reached confluency more rapidly than those cultured on PS. The reduction in time to confluency for SZCs on CM and FN may be the result of increased attachment, enhanced proliferation, or a combination of both. Increased attachment, as seen on CM and FN, results in a greater number of cells on the culture surfaces immediately following seeding, thus reducing the time required to achieve confluency. In addition, CM enhanced *CCND1* mRNA levels in SZC. CCND1 is a key regulator of cell cycle progression that promotes the transition from the G1 to the S phase of the cell cycle and is associated with proliferative activity [35]. The elevated *CCND1* expression observed on CM is therefore consistent with enhanced cell cycle progression and suggests that CM may promote proliferation. In contrast, FN reduced the time required for SZC to reach confluency despite no increases in *CCND1* mRNA levels, which suggests that *CCND1* is unlikely to account for the increase in proliferation on FN. Instead, the accelerated time to confluency on FN may reflect additional signaling pathways regulate proliferation independent of CCND1. Specifically, MRTF-A signaling has been implicated in regulating cell proliferation [12, 21, 36]. Therefore, FN may promote expansion through signaling pathways that differ from those operating on CM.

Although CM and FN similarly improved expansion, they had contrasting effects on the differentiation status of SZCs. Analysis of cell morphology highlighted the differences between FN and CM following SZC expansion. By P2, SZCs cultured on FN exhibited increased cell area and adopted a spread fibroblastic morphology similar to cells cultured on PS. In contrast, SZCs cultured on CM maintained a smaller cell area throughout passaging, consistent with the preservation of a more native SZC morphology. These findings suggest that CM contains additional matrix cues, beyond those provided by FN, that actively restrict cell spreading and SZC dedifferentiation. In support that SZCs were more dedifferentiated on FN, the changes in morphology are accompanied by differences in gene expression. FN did not prevent loss of chondrogenic or SZ-specific markers during passaging, while CM maintains a transcriptional profile relative to FN and PS. The prevention of dedifferentiation in CM could be due to at least three different factors. Firstly, unlike FN-coated tissue culture plastic, CM retains a complex mixture of cartilage-derived ECM proteins including Collagen Type VI (COLVI), Collagen Type XII (COLXII), and Collagen Type I (COL1) [26]. Among these, COLVI is of particular interest because it is a component of the chondrocyte pericellular matrix and has been implicated in maintaining chondrocyte phenotype [37, 38]. Similarly, culture on COL1 coated substrates has been shown to suppress chondrocyte dedifferentiation during monolayer expansion through maintaining a rounded morphology and reduced F-actin stress fibers [39]. Together, these findings suggest that the prevention of SZC dedifferentiation on CM may result from the combined actions of multiple matrix components that recapitulate the native extracellular environment. Secondly, surface topography generated by the deposited matrix by cells in CM could also play a role in limiting dedifferentiation as it has been shown that chondrocyte morphology and dedifferentiation is dependent on the microtopographies of culture surfaces [40]. Finally, in addition to topography, matrix stiffness is known to influence chondrocyte phenotype [41, 42]. CM matrices have a stiffness of ∼350kPa [43], which is order of magnitudes less than PS and may therefore prevent SZC dedifferentiation. These biochemical and biophysical features of CM may alter SZC behavior and delineating the contribution of each is worthy of future investigation.

The ability of CM, but not FN, to suppress SZC dedifferentiation suggested that these substrates differentially regulate actin organization and actin-mediated signaling. Expansion on PS and FN led to enhanced F-actin stress fiber formation and increased nuclear localization of MRTF-A. On the other hand, cells cultured on CM had less stress fibers and retained MRTF-A within the cytoplasm. MRTF-A is a mechanosensitive transcription factor whose activity is regulated by the balance between G- and F-actin. Increased actin polymerization promotes MRTF-A translocation into the nucleus, where it drives expression of fibroblastic and contractile genes [12, 21, 44]. Consistent with this mechanism, increased nuclear MRTF-A localization on PS and FN was accompanied by elevated expression of COL1, αSMA, and TNC, genes previously shown to be regulated by MRTF [12, 21]. Conversely, reduced stress fiber formation and predominantly cytoplasmic MRTF-A localization on CM were associated with suppression of these dedifferentiation markers. Collectively, these findings support a model in which the biochemical and biophysical characteristics of CM reduce SZC spreading, which prevents F-actin reorganization, suppresses MRTF-A nuclear translocation, leading to a reduction in fibroblastic matrix and contractile expression that is associated with chondrocyte dedifferentiation.

In conclusion, using bovine SZC, we demonstrate that FN alone is capable of promoting SZC attachment and proliferation, but does not suppress SZC dedifferentiation. Therefore, preserving the SZC phenotype during expansion may require a more complex extracellular matrix environment. Further studies that investigate the biochemical and biophysical characteristics implicit in CM would provide an additional understanding of the critical extracellular regulators of SZC phenotype. In addition, while the bovine SZCs in this study are obtained from generally healthy cartilage, generating a superior cell population for clinical therapeutic use will require a deeper understanding of SZCs that are derived from human cartilage and from cartilage tissues from different disease states. Defining the extracellular, matrix factors that regulate SZC attachment, proliferation and dedifferentiation will be instrumental for the generation of reparative tissue that consists of a native-like superficial zone.

## EXPERIMENTAL PROCEDURES

### Bovine Primary SZC Isolation

SZCs were isolated from the SZ of cartilage harvested from bovine metacarpal-phalangeal joints as previously described [10, 25]. Briefly, SZ cartilage was harvested from joints by microdissection and SZCs were extracted from tissue using sequential enzymatic digestion consisting of 0.5% protease (Sigma-Aldrich, 9036-06-0; St. Louis, MO) at 37°C for 45 minutes, followed by 0.1% collagenase (Roche, Mannheim, Germany) at 37°C for 14-17 hours. The resulting cell suspension was filtered, washed in phosphate buffered saline without calcium and magnesium (PBS), Quality Biological, Gaithersburg, MD, USA), and counted.

### Expansion on PS, CM, and FN

For monolayer expansion, SZCs were seeded on either PS (Corning, Edison, New Jersey, USA), FN-coated PS, or CM-coated PS. CM-coated PS dishes were obtained from a commercial source (StemBioSys, San Antonio, TX, USA). To coat PS with FN, FN (Fibronectin, Bovine Plasma, EMD Millipore Sigma, 341631; Darmstadt, Germany) at a concentration of 10 μg/mL in PBS was placed on top of PS dishes and maintained at room temperature for 2 hours. For coating a T75 flask, 10 mL of solution was added, and for an individual 6 well, 2 mL of solution were added for a final concentration of 2.5 μg/cm^2^. After 2 hours, the surface was washed 3 times with PBS before seeding chondrocytes.

SZC were seeded at a density of 1500 cells/cm^2^ on PS, CM, and FN. Chondrocytes were maintained and passaged in expansion media. Expansion media consisted of Hams F-12 media (Corning, MT10080CV) supplemented with 10% FBS (GenClone, Genesee Scientific, San Diego, CA, USA) and 1% antibiotic-antimycotic (Corning). SZCs were cultured until approximately 80% to 90% confluent, then detached using 0.25% trypsin-EDTA (GenClone). Following detachment cells were deemed as passage 1 (P1) cells. P1 SZCs from each condition (PS, CM, FN) were then reseeded in monolayer (2 × 10^3^ cells/cm^2^) on their respective surfaces. P1 cells were then expanded until ∼80%-90% confluency and then detached with 0.25% trypsin-EDTA to obtain passage 2 (P2) cells.

### Attachment and Confluency Analysis

To quantify cell area coverage (confluency) and attachment, P0 SZC were seeded in monolayer at 5.0 × 10^4^ cells/cm^2^ on either PS, CM, or FN in an expansion medium. After 24 hours, nonadherent cells were removed by aspirating the media and washing the cultures with PBS. Light microscopy images were obtained using a Zeiss Primovert tissue culture microscope (Zeiss, Jena, Germany) equipped with an attached camera (Swiftcam Technologies; Hong Kong, China). Images were imported into FIJI (ImageJ; National Institutes of Health) for analysis. Adherent cells within each field were manually traced and outlined using the freehand selection tool. The total traced cell area was calculated and consequently divided by the total image area to determine the percent area covered. Multiple fields were quantified per well, with a minimum of 3 wells analyzed per experimental condition. For assessing confluency throughout passaging, this procedure was performed on each of the SZC cultures per day throughout passaging from P0 to P1 and from P1 to P2, allowing cell expansion to be compared over time across conditions.

The proportion of cells that attached onto each culture surface was quantified by DNA quantification as previously described [10, 26]. Briefly, P0 SZC were seeded at a density of 5.0 × 10^4^ cells/cm^2^ onto either PS, CM, or FN in monolayer expansion media. After 24 hours, non-adherent cells were removed by aspirating the media, and the cells were gently washed with PBS. Adherent cells were harvested using a papain digestion solution (40 μg/ml papain in 20 mM ammonium acetate, 1 mM EDTA, and 2 mM dithiothreitol at pH 6.2), followed by digestion for 4 hours at 65°C. DNA content from adherent cells on each culture surface was compared with the DNA content of aliquots taken from the initial cell suspension at the time of seeding to determine the proportion of cells that attached. Quantification of DNA content from papain digested cell pellets was performed using the Hoechst 33258 dye-binding assay. A standard curve was generated with calf thymus DNA diluted in PBS. To measure absorbance, fluorescent spectrophotometry (Glomax Multi+ Detection System; excitation, λ = 355 nm; emission, λ = 460 nm) was used.

### RNA Extraction and Relative RT-PCR

Total RNA was isolated from SZCs using TRIzol reagent (Sigma-Aldrich) in accordance with the manufacturer’s instructions. RNA yield and purity were evaluated with a NanoDrop One spectrophotometer (Thermo Fisher Scientific, Waltham, MA, USA). RNA was reversed transcribed into complementary DNA (cDNA) using UltraScript 2.0 cDNA Synthesis Kit (PCR Biosystems; Wayne, PA, USA), following the manufacturer’s protocols. Relative RT-PCR was performed using 20 ng of cDNA per reaction, previously validated gene-specific primers at 10 μM working concentration (sequences, GenBank accession numbers, and temperature provided in 01) [11, 45], and qPCRBIO SyGreen Blue Master Mix (PCR Biosystems).

Reactions were conducted in duplicate on a Cielo 3 Real-Time PCR System (Azure Biosystems, Houston, TX, USA; see 01 for specific PCR conditions). No-template controls and no reverse transcriptase controls were included in each run to verify specificity and exclude contamination. Mean relative quantification values were calculated using the Pfaffl method with 18S rRNA as the endogenous reference gene [46].

### Morphology, Area, and Circularity

To assess morphology, SZCs at P0 and P2 were seeded in monolayer at 5.0 × 10^4^ cells/cm^2^ on PS, CM, and FN in expansion media. After 24 hours, light microscopy images were obtained using a Zeiss Primovert tissue culture microscope equipped with a camera (Swiftcam). Cell boundaries of adherent cells were manually traced in FIJI. Cell area and circularity were calculated as previously described [22, 26]. Circularity values approaching 1 represent more rounded cells, whereas lower values reflect increasingly elongated, fibroblastic morphologies. At least 3 image fields were evaluated per condition, with a minimum of 45 cells analyzed per condition.

### Immunofluorescent Staining of αSMA and MRTF-A

To examine αSMA positive stress fibers and MRTF-A localization, P2 SZCs expanded on PS, CM and FN were seeded at 1.0 × 10^4^ cells/cm^2^ on non-coated, CM-coated Thermanox Plastic coverslips, or FN-coated Thermanox Plastic coverslips (Thermofisher). After 24 hours, cells were fixed by incubating cells in 4% paraformaldehyde (Electron Microscopy Sciences, Hatfield, PA, USA) for 15 minutes at room temperature, followed by three washes with PBS. Cells were then permeabilized and blocked using a solution containing 3% Goat serum, 3% BSA, and 0.3% Triton for 30 minutes. Following permeabilization, cells were incubated overnight at 4°C with primary antibody against αSMA (Mouse Monoclonal, Dilution 1:200, ab7817, ABCAM Cambridge, United Kingdom) or with a primary antibody against MRTF-A (Rabbit Monoclonal, Dilution 1:100, 77098, Cell Signaling Technologies, Danvers, MA, USA). After 24 hours, cells were washed and incubated for 1 hour at room temperature with secondary antibody solution consisting of Hoechst 33342 (Nuclei; 1/500; Biotium, 40046; Fremont, CA), Rhodamine phalloidin (F-actin; 1:50; Biotium, 00027; Fremont, CA), and CF488A Goat Anti-Rabbit IgG (MRTF; 1:500, Biotium, 20019) or CF488A Goat Anti-Mouse IgG (αSMA; 1:500, Biotium, 20302). Cells were washed three times with PBS between incubations. Slides were then mounted with Drop-n-Stain mounting medium (Biotium) and glass coverslips. Confocal microscopy was performed using a Zeiss LSM880 laser scanning confocal fluorescence microscope with a 40x objective. Z-stack images were acquired using 0.4-μm slices.

### Quantification of αSMA Positive Stress Fibers

The proportion of cells that were positive for αSMA in stress fibers were quantified in FIJI by manually assessing individual cells for the presence of organized αSMA-positive stress fibers, defined as distinct bundled filamentous structures spanning the cell body. For each condition within a single experimental set, four images were analyzed, and 25 cells were evaluated per image, yielding 100 cells per condition. The percentage of αSMA positive cells was calculated for each condition by dividing the number of positive cells by the total cells counted. This procedure was repeated across three independent experimental sets, totaling 300 cells evaluated per condition.

### Analysis of MRTF-A Nuclear Localization

For quantitative analysis, MRTF-A nuclear and cytoplasmic fluorescence intensity was measured in FIJI as previously described [47]. Nuclear regions were defined using Hoechst signal, and cell boundaries were identified using phalloidin labeled F-actin. Integrated MRTF-A fluorescence within nuclear and cytoplasmic regions was used to calculate the nuclear/cytoplasmic MRTF-A ratio for each cell.

### Statistical Analysis

All experiments were conducted using SZCs isolated from multiple bovine joints and were repeated in at least three independent runs unless otherwise stated. Each independent run represents a separate SZC isolation performed on a different day using cells derived from different bovine donors. For analysis, data from replicate experiments were consolidated. Statistical analyses were performed using GraphPad Prism (GraphPad Software Inc, Boston, MA). Data sets were screened for outliers using the ROUT method [48]. Comparisons among PS, CM, and FN groups were performed using one-way analysis of variance (ANOVA) followed by appropriate multiple-comparisons testing. Data are reported as mean ± standard deviation, and statistical significance was defined by a predetermined alpha level.

**Table 1:**
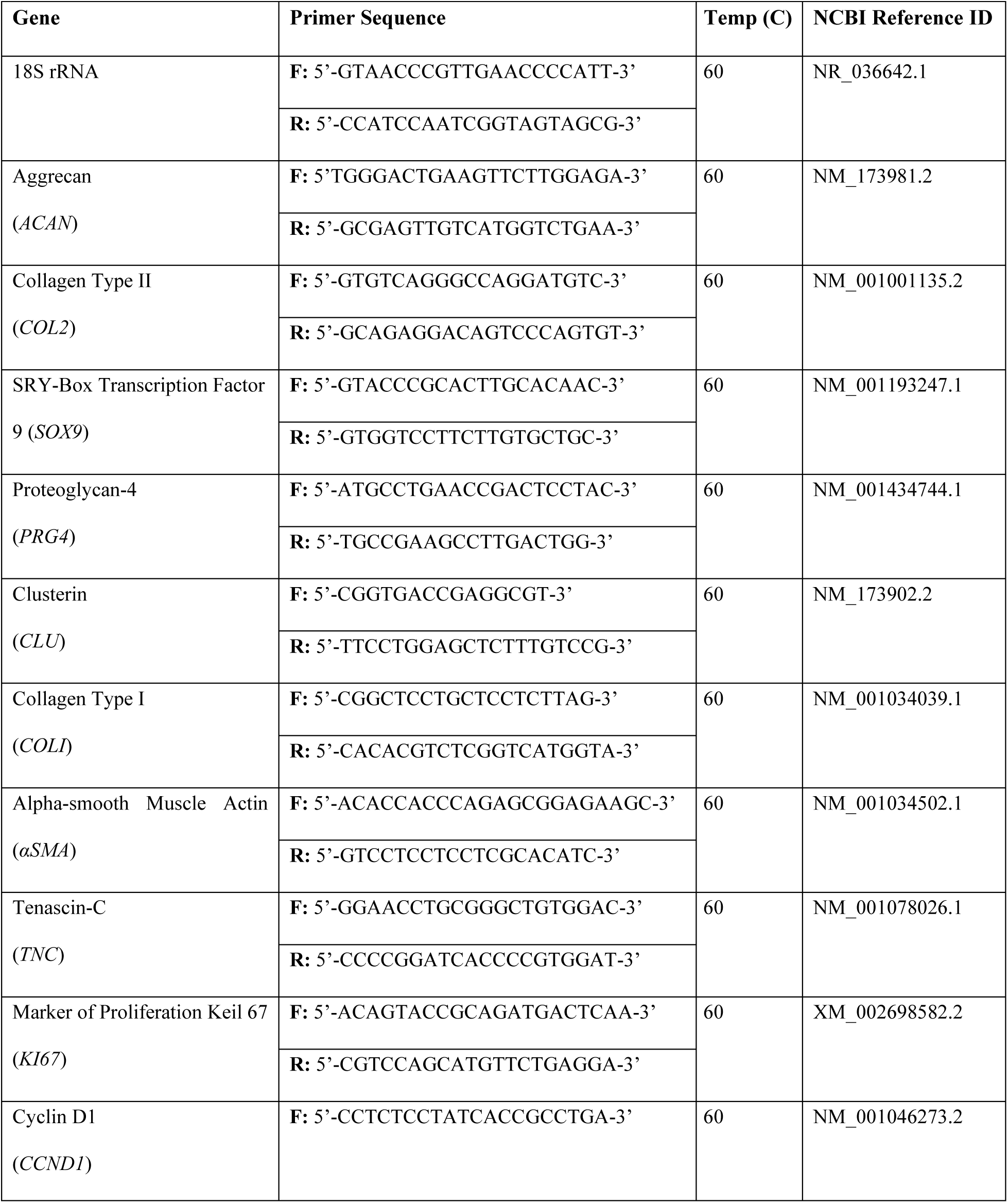
Gene-specific primer sequences.

## ACKNOWLEDGEMENTS

This work was supported in part by the Orthopaedic Research and Education Foundation (OREF, 26-010, awarded to AWS and JP).

## REFERENCES

[1] C.K. Jung, Articular Cartilage: Histology and Physiology, in: S.-J.K. Asode Ananthram Shetty, Norimasa Nakamura,Mats Brittberg (Ed.), Techniques in Cartilage Repair Surgery, Springer Heidelberg New York Dordrecht London2014, pp. 17–22.

[2] S. Fox, A. J., A. Bedi, S.A. Rodeo, The basic science of articular cartilage: structure, composition, and function, Sports Health 1(6) (2009) 461–8.

[3] C.R. Flannery, C.E. Hughes, B.L. Schumacher, D. Tudor, M.B. Aydelotte, K.E. Kuettner, B. Caterson, Articular cartilage superficial zone protein (SZP) is homologous to megakaryocyte stimulating factor precursor and Is a multifunctional proteoglycan with potential growth-promoting, cytoprotective, and lubricating properties in cartilage metabolism, Biochem Biophys Res Commun 254(3) (1999) 535–41.

[4] M.Z. Ruan, A. Erez, K. Guse, B. Dawson, T. Bertin, Y. Chen, M.M. Jiang, J. Yustein, F. Gannon, B.H. Lee, Proteoglycan 4 expression protects against the development of osteoarthritis, Sci Transl Med 5(176) (2013) 176ra34.

[5] M. Brittberg, A. Lindahl, A. Nilsson, C. Ohlsson, O. Isaksson, L. Peterson, Treatment of deep cartilage defects in the knee with autologous chondrocyte transplantation, N Engl J Med 331(14) (1994) 889–95.

[6] D.A. Grande, M.I. Pitman, L. Peterson, D. Menche, M. Klein, The repair of experimentally produced defects in rabbit articular cartilage by autologous chondrocyte transplantation, J Orthop Res 7(2) (1989) 208–18.

[7] E.A. Makris, A.H. Gomoll, K.N. Malizos, J.C. Hu, K.A. Athanasiou, Repair and tissue engineering techniques for articular cartilage, Nat Rev Rheumatol 11(1) (2015) 21–34.

[8] L. Peterson, M. Brittberg, I. Kiviranta, E.L. Akerlund, A. Lindahl, Autologous chondrocyte transplantation. Biomechanics and long-term durability, Am J Sports Med 30(1) (2002) 2–12.

[9] M. Siczkowski, F.M. Watt, Subpopulations of chondrocytes from different zones of pig articular cartilage. Isolation, growth and proteoglycan synthesis in culture, J Cell Sci 97 (Pt 2) (1990) 349–60.

[10] E.E.R. Davis, T.J. Manzoni, V.J. Bianchi, J.F. Weber, P.H. Wu, S.C. Regmi, S.D. Waldman, T.A. Schmidt, A.W. Su, R.A. Kandel, J. Parreno, Passaged Articular Chondrocytes From the Superficial Zone and Deep Zone Can Regain Zone-Specific Properties After Redifferentiation, Am J Sports Med (2024) 3635465241230031.

[11] J. Parreno, M. Nabavi Niaki, K. Andrejevic, A. Jiang, P.H. Wu, R.A. Kandel, Interplay between cytoskeletal polymerization and the chondrogenic phenotype in chondrocytes passaged in monolayer culture, J Anat 230(2) (2017) 234–248.

[12] J. Parreno, S. Raju, P.H. Wu, R.A. Kandel, MRTF-A signaling regulates the acquisition of the contractile phenotype in dedifferentiated chondrocytes, Matrix Biol 62 (2017) 3–14.

[13] P.D. Benya, J.D. Shaffer, Dedifferentiated chondrocytes reexpress the differentiated collagen phenotype when cultured in agarose gels, Cell 30(1) (1982) 215–24.

[14] H. Holtzer, J. Abbott, J. Lash, S. Holtzer, The Loss of Phenotypic Traits by Differentiated Cells in Vitro, I. Dedifferentiation of Cartilage Cells, Proc Natl Acad Sci U S A 46(12) (1960) 1533–42.

[15] F. Mallein-Gerin, R. Garrone, M. van der Rest, Proteoglycan and collagen synthesis are correlated with actin organization in dedifferentiating chondrocytes, Eur J Cell Biol 56(2) (1991) 364–73.

[16] S. Roberts, J. Menage, L.J. Sandell, E.H. Evans, J.B. Richardson, Immunohistochemical study of collagen types I and II and procollagen IIA in human cartilage repair tissue following autologous chondrocyte implantation, Knee 16(5) (2009) 398–404.

[17] R.F. LaPrade, L.S. Bursch, E.J. Olson, V. Havlas, C.S. Carlson, Histologic and immunohistochemical characteristics of failed articular cartilage resurfacing procedures for osteochondritis of the knee: a case series, Am J Sports Med 36(2) (2008) 360–8.

[18] B. Kinner, M. Spector, Smooth muscle actin expression by human articular chondrocytes and their contraction of a collagen-glycosaminoglycan matrix in vitro, J Orthop Res 19(2) (2001) 233–41.

[19] A.T. Rzepski, M.M. Schofield, S. Richardson-Solorzano, M.L. Arranguez, A.W. Su, J. Parreno, Targeting the reorganization of F-actin for cell-based implantation cartilage repair therapies, Differentiation 143 (2025) 100847.

[20] S. Gonzalez-Nolde, C.J. Schweiger, E.E.R. Davis, T.J. Manzoni, S.M.I. Hussein, T.A. Schmidt, S.G. Cone, G.D. Jay, J. Parreno, The Actin Cytoskeleton as a Regulator of Proteoglycan 4, Cartilage (2024) 19476035231223455.

[21] J. Parreno, S. Raju, M.N. Niaki, K. Andrejevic, A. Jiang, E. Delve, R. Kandel, Expression of type I collagen and tenascin C is regulated by actin polymerization through MRTF in dedifferentiated chondrocytes, FEBS Lett 588(20) (2014) 3677–84.

[22] M.M. Schofield, A.T. Rzepski, S. Richardson-Solorzano, J. Hammerstedt, S. Shah, C.E. Mirack, M. Herrick, J. Parreno, Targeting F-actin stress fibers to suppress the dedifferentiated phenotype in chondrocytes, Eur J Cell Biol 103(2) (2024) 151424.

[23] M.B. Asparuhova, J. Ferralli, M. Chiquet, R. Chiquet-Ehrismann, The transcriptional regulator megakaryoblastic leukemia-1 mediates serum response factor-independent activation of tenascin-C transcription by mechanical stress, FASEB J 25(10) (2011) 3477–88.

[24] L.L. Luchsinger, C.A. Patenaude, B.D. Smith, M.D. Layne, Myocardin-related transcription factor-A complexes activate type I collagen expression in lung fibroblasts, J Biol Chem 286(51) (2011) 44116–44125.

[25] E. Delve, V. Co, R.A. Kandel, Superficial and deep zone articular chondrocytes exhibit differences in actin polymerization status and actin-associated molecules in vitro, Osteoarthr Cartil Open 2(3) (2020) 100071.

[26] T.J. Manzoni, A. Ho, L. Smull, V.C. West, J.D.V. Waters, K. Lemus, J. Adams, A.W. Su, J. Parreno, Enhanced Superficial Zone Chondrocyte Expansion and Redifferentiation by Culture on Chondrocyte-Derived Decellularized Matrices, Cartilage (2025) 19476035251369735.

[27] R.F. Loeser, Integrins and chondrocyte-matrix interactions in articular cartilage, Matrix Biol 39 (2014) 11–6.

[28] T. Tao, Y. Li, C. Gui, Y. Ma, Y. Ge, H. Dai, K. Zhang, J. Du, Y. Guo, Y. Jiang, J. Gui, Fibronectin Enhances Cartilage Repair by Activating Progenitor Cells Through Integrin alpha5beta1 Receptor, Tissue Eng Part A 24(13-14) (2018) 1112–1124.

[29] M. Uetaki, N. Onishi, Y. Oki, T. Shimizu, E. Sugihara, O. Sampetrean, T. Watanabe, H. Yanagi, K. Suda, H. Fujii, K. Kano, H. Saya, H. Nobusue, Regulatory roles of fibronectin and integrin alpha5 in reorganization of the actin cytoskeleton and completion of adipogenesis, Mol Biol Cell 33(9) (2022) ar78.

[30] V. Lefebvre, W. Huang, V.R. Harley, P.N. Goodfellow, B. de Crombrugghe, SOX9 is a potent activator of the chondrocyte-specific enhancer of the pro alpha1(II) collagen gene, Mol Cell Biol 17(4) (1997) 2336–46.

[31] Q. Zhang, Q. Ji, X. Wang, L. Kang, Y. Fu, Y. Yin, Z. Li, Y. Liu, X. Xu, Y. Wang, SOX9 is a regulator of ADAMTSs-induced cartilage degeneration at the early stage of human osteoarthritis, Osteoarthritis Cartilage 23(12) (2015) 2259–2268.

[32] G.P. Dowthwaite, J.C. Bishop, S.N. Redman, I.M. Khan, P. Rooney, D.J. Evans, L. Haughton, Z. Bayram, S. Boyer, B. Thomson, M.S. Wolfe, C.W. Archer, The surface of articular cartilage contains a progenitor cell population, J Cell Sci 117(Pt 6) (2004) 889–97.

[33] R.O. Hynes, Integrins: versatility, modulation, and signaling in cell adhesion, Cell 69(1) (1992) 11–25.

[34] C.M. West, H. de Weerd, K. Dowdy, A. de la Paz, A specificity for cellular fibronectin in its effect on cultured chondroblasts, Differentiation 27(1) (1984) 67–73.

[35] B. Ma, J.C. Leijten, L. Wu, M. Kip, C.A. van Blitterswijk, J.N. Post, M. Karperien, Gene expression profiling of dedifferentiated human articular chondrocytes in monolayer culture, Osteoarthritis Cartilage 21(4) (2013) 599–603.

[36] A. Wells, A. Nuschke, C.C. Yates, Skin tissue repair: Matrix microenvironmental influences, Matrix Biol 49 (2016) 25–36.

[37] R. Quarto, B. Dozin, P. Bonaldo, R. Cancedda, A. Colombatti, Type VI collagen expression is upregulated in the early events of chondrocyte differentiation, Development 117(1) (1993) 245–51.

[38] J. Gouttenoire, U. Valcourt, M.C. Ronziere, E. Aubert-Foucher, F. Mallein-Gerin, D. Herbage, Modulation of collagen synthesis in normal and osteoarthritic cartilage, Biorheology 41(3-4) (2004) 535–42.

[39] M. Kino-Oka, S. Yashiki, Y. Ota, Y. Mushiaki, K. Sugawara, T. Yamamoto, T. Takezawa, M. Taya, Subculture of chondrocytes on a collagen type I-coated substrate with suppressed cellular dedifferentiation, Tissue Eng 11(3-4) (2005) 597–608.

[40] E. Costa Martinez, J.C. Rodriguez Hernandez, M. Machado, J.F. Mano, J.L. Gomez Ribelles, M. Monleon Pradas, M. Salmeron Sanchez, Human chondrocyte morphology, its dedifferentiation, and fibronectin conformation on different PLLA microtopographies, Tissue Eng Part A 14(10) (2008) 1751–62.

[41] N.L. Bergholt, M. Foss, A. Saeed, N. Gadegaard, H. Lysdahl, M. Lind, C.B. Foldager, Surface chemistry, substrate, and topography guide the behavior of human articular chondrocytes cultured in vitro, J Biomed Mater Res A 106(11) (2018) 2805–2816.

[42] W. Zhong, Y. Li, L. Li, W. Zhang, S. Wang, X. Zheng, YAP-mediated regulation of the chondrogenic phenotype in response to matrix elasticity, J Mol Histol 44(5) (2013) 587–95.

[43] Y. Mao, T. Block, A. Singh-Varma, A. Sheldrake, R. Leeth, S. Griffey, J. Kohn, Extracellular matrix derived from chondrocytes promotes rapid expansion of human primary chondrocytes in vitro with reduced dedifferentiation, Acta Biomater 85 (2019) 75–83.

[44] E. Langelier, R. Suetterlin, C.D. Hoemann, U. Aebi, M.D. Buschmann, The chondrocyte cytoskeleton in mature articular cartilage: structure and distribution of actin, tubulin, and vimentin filaments, J Histochem Cytochem 48(10) (2000) 1307–20.

[45] E. Delve, J. Parreno, V. Co, P.H. Wu, J. Chong, M. Di Scipio, R.A. Kandel, CDC42 regulates the expression of superficial zone molecules in part through the actin cytoskeleton and myocardin-related transcription factor-A, J Orthop Res 36(9) (2018) 2421–2430.

[46] M.W. Pfaffl, A new mathematical model for relative quantification in real-time RT-PCR, Nucleic Acids Res 29(9) (2001) e45.

[47] V.C. West, K.E. Owen, K.L. Inguito, K.M.M. Ebron, T.N. Reiner, C.E. Mirack, C.H. Le, R. de Cassia Marqueti, S. Snipes, R. Mousavizadeh, R.E. King, D.M. Elliott, J. Parreno, Actin Polymerization Status Regulates Tenocyte Homeostasis Through Myocardin-Related Transcription Factor-A, Cytoskeleton (Hoboken) (2024).

[48] H.J. Motulsky, R.E. Brown, Detecting outliers when fitting data with nonlinear regression - a new method based on robust nonlinear regression and the false discovery rate, BMC Bioinformatics 7 (2006) 123.

